# Allocating maternal-tree collections to conserve genetic diversity of *Lumnitzera littorea* in southern Vietnam

**DOI:** 10.64898/2026.11.26.769528

**Authors:** Duc-Hoan Huynh, The-Kiet Bui-Nguyen, Thanh-Cong Nguyen, Hoang-Dung Tran

**Author notes:** Correspondence to Duc-Hoan Huynh.

## Abstract

Mangrove restoration nurseries often raise seed from a limited number of maternal trees, and how those trees are split among source populations affects which alleles reach the nursery. We genotyped 110 *Lumnitzera littorea* trees from two mainland and two island sites in southern Vietnam at 11 nuclear microsatellite loci and then enumerated all 5,236 ways of dividing a 30-tree collection among the four sites. With sample sizes matched, the mainland sites had consistently higher expected heterozygosity than the islands. Con Dao and Phu Quoc were the most differentiated pair on the full panel; dropping two influential loci put Can Gio and Phu Quoc at the top instead. Neither pattern changed when we compared the primary genotype calls with an earlier set of calls, restored 23 calls from overlapping adjacent peaks, or removed loci with high estimated null-allele frequencies. Taking 6, 8, 11, and 5 trees from Can Gio, Dong Nai, Con Dao, and Phu Quoc retained an expected 36.89 of 37 alleles on the full panel and 29.94 of 30 on the reduced panel. A nearly even split of 8, 7, 8, and 7 trees gave Phu Quoc better representation and cost little in pooled allele capture. Either split keeps all four provenances in a small collection; which one to use depends on whether a program values the site-private alleles at Con Dao more than balanced representation of every source.

## 2 Introduction

Restoring fragmented mangrove forests depends partly on how seed is collected. Time, site access, and seed availability often cap the number of maternal trees, and the obvious response, collecting where trees are plentiful, can leave other sources barely represented. The planning question is how much neutral diversity a fixed number of trees can capture while every provenance is still included. Putting numbers on that trade-off helps both in designing seed collections and in documenting the source material used in restoration (Nef et al., 2021).

*Lumnitzera littorea* (Combretaceae) occurs across the Indo-West Pacific and is locally threatened. In southern Vietnam it survives at mainland and island sites that are managed under different conditions (Su et al., 2007; Huynh et al., 2026). Our previous survey of these sites with inter-simple sequence repeats (ISSRs) found substantial variation among them (Huynh et al., 2026), but binary band scores describe band variation and population differentiation, not which tree carries which allele. Codominant microsatellites fill that gap: they give observed heterozygosity and, when the electropherogram supports the call, a tree-by-tree record of alleles, which is what a model of maternal-tree collections needs. Guo et al. (2021) used the same microsatellite panel to describe differentiation across the species^’^ range.

We first asked whether the diversity ranking of the four sites, and their pairwise differentiation, depended on how alleles were scored and which loci were used. We then asked how well different allocations of 30 maternal trees would retain the alleles we observed, judged in three ways: pooled allele capture, representation within each provenance, and the chance of keeping alleles found at only one site. The three criteria do not always agree. An allocation can capture almost every pooled allele and still leave a source unrepresented.

## 3 Materials and methods

### 3.1 Sampling and scope of the dataset

We analyzed 110 *L. littorea* trees from Can Gio (30), Dong Nai (30), Con Dao (20), and Phu Quoc (30). The mainland trees genotyped with SSRs were the same trees analyzed with ISSRs from the same field campaign, so we assigned them to sites using the ISSR sample order: trees C1–C30 were from Can Gio and C31–C60 from Dong Nai. Sites and sampling areas are described in Section 4.1 and Figure 3 of Huynh et al. (2026). Five *L. racemosa* samples were included only in the ordination as interspecific references; all population analyses and seed-collection models used *L. littorea* alone.

### 3.2 DNA extraction and microsatellite amplification

Total DNA was isolated from dried leaf tissue using the cetyltrimethylammonium bromide (CTAB) method (Doyle and Doyle, 1990), with the modifications reported by Huynh et al. (2026). Approximately 100 mg of dry leaf tissue was ground in liquid nitrogen, mixed with 700 µL of preheated 2% CTAB buffer, and incubated at 65 °C for 30 min. The extraction buffer contained 100 mM Tris-HCl (pH 8.0), 20 mM EDTA, 1.4 M NaCl, 1% PVP-40, and 0.2% β-mercaptoethanol. After extraction with chloroform and isoamyl alcohol (24:1), DNA was precipitated in cold isopropanol, washed with 70% ethanol, and dissolved in 50 µL of TE buffer. DNA quality was checked on 0.8% agarose in 1× TBE buffer after staining with 1× SYBR Safe for 10–15 min and visualization under a 470-nm blue LED.

The 11 nuclear microsatellites were LL-144, LL-379, LL-852, LL-1086, LL-1150, LL-2749, LL-2952, LL-3258, LL-3573, LL-4058, and LL-4164 (Guo et al., 2021). Following the SSR amplification protocol of Guo et al. (2021), each 20-µL PCR contained 20 ng of template DNA, 1× PCR buffer, 1 mM dNTPs, 1.0 µL of each primer, and 1 U of Taq DNA polymerase. Cycling consisted of an initial denaturation at 94 °C for 4 min; 30 cycles of 94 °C for 40 s, 53 °C for 1 min, and 72 °C for 1 min; and a final extension at 72 °C for 10 min. Primer sequences are provided in Supplementary Dataset S4. PCR fragments were analyzed by capillary electrophoresis on the BiOptic Qsep platform. The reports list program M-4-10-06-300, which specifies injection at 4 kV for 10 s and separation at 6 kV for 300 s (BiOptic Inc., 2023, pp. 32–33). The 20–1K alignment marker contained 20-bp and 1,000-bp reference fragments. The reports provided fragment size, fluorescence intensity in relative fluorescence units (RFU), and peak integration intervals.

### 3.3 Allele calling and sensitivity analyses

Peaks were assigned to locus-specific size bins. At LL-852, LL-2749, LL-2952, LL-3258, and LL-3573, the secondary-peak threshold was 15–30% of the main peak, depending on the locus. For the other six loci (LL-144, LL-379, LL-1086, LL-1150, LL-4058, and LL-4164), the technical reports specified a 10 RFU minimum. At these loci, the main peak had to reach 10 RFU, and a candidate second peak had to fall in a different bin, reach both 10 RFU and 40% of the main peak, and be distinguishable from it. At LL-4164, a peak 3 bp shorter than the main peak was flagged as possible stutter (Van Oosterhout et al., 2004; Hoffman and Amos, 2005).

Initial scoring left 163 tree–locus combinations unresolved. We reviewed the original and repeat reports for all 163, including the 148 combinations re-amplified in the final repeat-PCR round. We called two alleles when both fragment sizes recurred and the peaks met the bin, intensity, separation, and stutter criteria. In 23 combinations, the two reported maxima were 1 bp apart and their integration intervals overlapped in all 47 traces examined. The overlap prevented us from confirming peak separation from these reports. We therefore treated all 23 combinations as unresolved in the primary matrix (40% cutoff) and in the 30% sensitivity matrix. Supplementary Datasets S1–S3 document the scoring history and sources, and Supplementary Dataset S8 compares the primary results with a matrix that retains these 23 calls.

### 3.4 Diversity and differentiation

For each site and locus, we calculated observed heterozygosity (*H*_*O*_) and expected heterozygosity (*H*_*E*_) from the available genotypes. Allelic richness (*A*_*R*_) was rarefied to 22 allele copies per site and locus so that the earlier, primary, and 30% threshold matrices could be compared on a common basis (Kalinowski, 2004). We obtained multilocus *F*_*IS*_ from summed within-site variance components (Weir and Cockerham, 1984). To compare *H*_*E*_ at a common sample size, we subsampled each site to the smallest valid sample size at each locus, without replacement, in 12,000 replicates and applied the finite-sample correction before averaging across loci. To test whether unresolved calls were concentrated at particular sites, we permuted site labels among trees 19,999 times and applied an exact, fixed-margin test to each locus, with Holm adjustment of the locus-level tests.

We assessed Hardy–Weinberg proportions separately for each polymorphic site–locus combination using 49,999 random pairings of allele copies conditional on their observed counts (Engels, 2009). The two-sided and one-sided test probabilities were adjusted separately with the Benjamini– Hochberg procedure. Null-allele frequencies were estimated for each site–locus combination with the first estimator of Brookfield (1996), and the diversity and differentiation analyses were repeated after excluding loci with an estimate of 0.20 or higher at any site. We estimated pairwise and overall *F*_*ST*_ with the method of Weir and Cockerham (1984) and calculated percentile intervals from 20,000 bootstrap resamples of loci. Hierarchical analysis of molecular variance partitioned variation among the mainland and island groups, among sites within groups, and within sites (Excoffier et al., 1992). For the group-level test, the mainland–island grouping was compared with all three ways of dividing the four sites into two pairs, so the smallest attainable probability was 0.333.

To examine the effect of marker composition, we compared the full 11-locus panel with a nine-locus panel that omitted LL-4058 and LL-4164, which influenced the between-island estimate. The five loci without unresolved calls were used for a principal coordinate analysis of the square-root allele-sharing distance, first for *L. littorea* and then with the five congener samples. We compared patterns with the published ISSR results for the same localities (Huynh et al., 2026). Published site-level estimates from Guo et al. (2021), including expected heterozygosity from comparably sized samples, provided regional context; for descriptive comparison with their allelic richness table, we also calculated richness at 14 allele copies.

### 3.5 Maternal-tree allocation and allele capture

We enumerated all feasible allocations of 20, 30, 40, and 50 trees across the four sites, sampling trees without replacement within each site; the 30-tree comparison served as the main planning example. If a site contains N sampled trees and m of them carry an allele, the probability of missing the allele when drawing n trees is C(N − m, n)/C(N, n). We multiplied these probabilities across sites to obtain the probability of missing that allele and summed the complementary probabilities over the observed alleles to estimate pooled capture. For unresolved calls outside the 23 adjacent-peak cases, the reported main-peak bin was used in the pooled-capture model. The 23 cases contributed neither allele in the primary and 30% matrices. We used the inclusion–exclusion principle to estimate the joint probability of retaining all site-private alleles. Within-site representation was the expected number of that site^’^s observed alleles represented by trees drawn from that site. All of these quantities refer to alleles carried by the sampled maternal trees.

We compared three kinds of allocation: unconstrained, at least five trees per source, and nearly even, with site counts differing by no more than one tree. For each scoring matrix and panel, expected pooled capture was expressed relative to the best value possible with at least five trees per source. Each allocation was then scored by its lowest relative capture across matrices and panels, and we picked the highest-scoring allocation among those with at least five trees per source and among the nearly even ones. The supplementary script reproduces the allocation search.

## 4 Results

### 4.1 Allele-call sensitivity and genetic diversity

The repeat review initially produced two-allele calls for 154 of the 163 unresolved combinations. After the 23 adjacent-peak pairs were set to unresolved, 131 two-allele calls remained, 117 of them among the 148 combinations in the final repeat-PCR round. The primary matrix therefore contained 32 unresolved entries among 1,210 *L. littorea* genotypes (2.64%): 9, 9, 7, and 7 at Can Gio, Dong Nai, Con Dao, and Phu Quoc, respectively (Supplementary Table S1). Missing calls showed no detectable concentration by site (permutation *p* = 0.879). Five of the originally unresolved cases at LL-4164 kept a possible-stutter flag, and four fell just below the 40% threshold. A further signal at 244–247 bp at LL-144 fell outside the 198–200 bp scoring interval and was not scored.

Expected heterozygosity was higher at the two mainland sites than at either island on both panels (Table 1, Figure 1). The sample-size-matched 11-locus means were 0.524, 0.554, 0.340, and 0.296 for Can Gio, Dong Nai, Con Dao, and Phu Quoc, respectively, and both mainland sites exceeded both islands in every 11-locus resampling replicate under all three scoring matrices. Between the earlier and primary calls, *H*_*O*_ rose from 0.116 to 0.333 at Dong Nai and from 0.102 to 0.217 at Can Gio, and *F*_*IS*_ fell from 0.795 to 0.406 and from 0.808 to 0.592, respectively. In the primary matrix, 24 of 37 polymorphic site–locus combinations departed from Hardy–Weinberg proportions at *q* < 0.05, and 17 showed heterozygote deficiency in the one-sided test. Under the 30% rule, the corresponding counts were 23 and 17. Brookfield estimates reached 0.20 or more in nine site–locus combinations, all at LL-852, LL-1086, LL-2952, or LL-3573 (Supplementary Dataset S7), and none of the 60 mainland trees was scored as heterozygous at LL-2952. Without these four loci, both mainland sites still had higher *H*_*E*_ than both islands in every resampling replicate, and *F*_*IS*_ fell to 0.434 at Can Gio and 0.280 at Dong Nai (Supplementary Dataset S6).

**Table 1.**
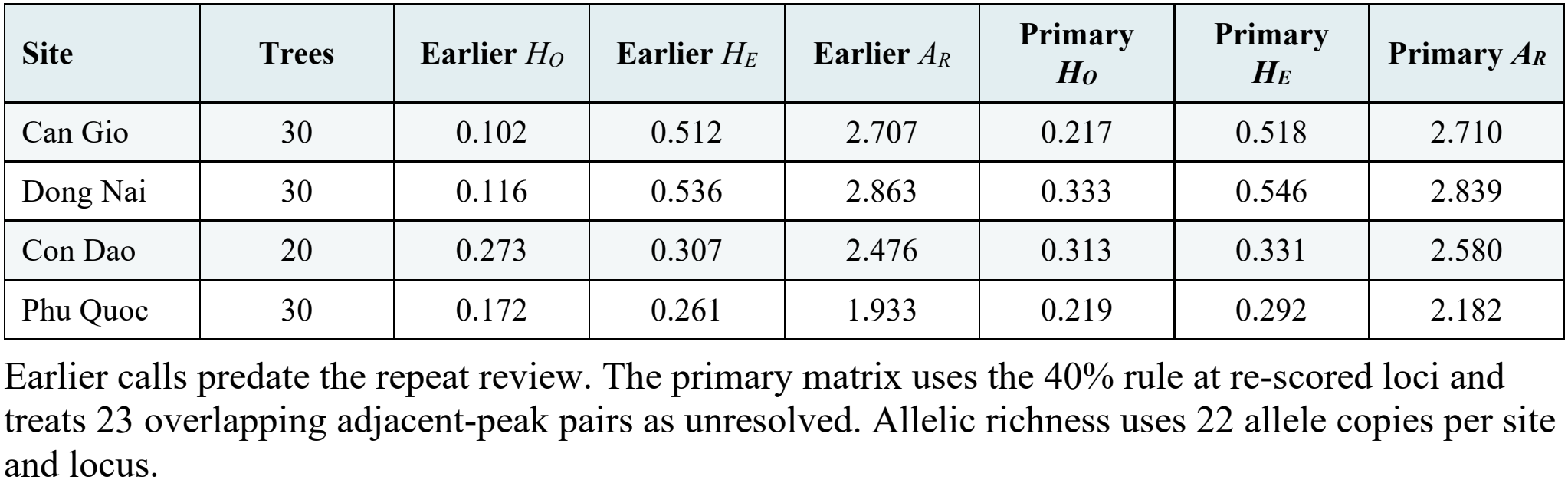
Diversity at the 11 nuclear microsatellite loci.

**Figure 1.**
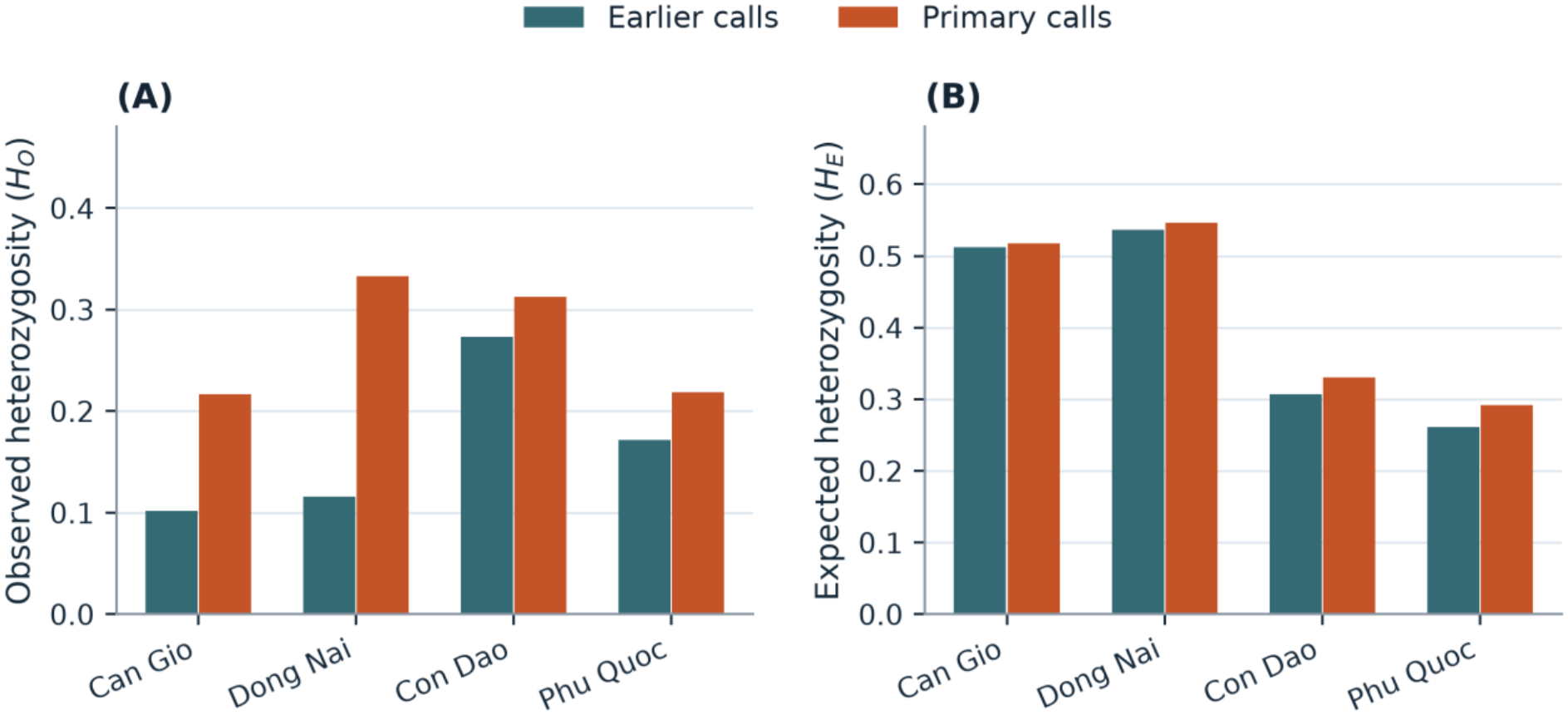
Observed (A) and expected (B) heterozygosity at the four sites on the 11-locus panel. Bars compare earlier calls with primary calls.

### 4.2 Differentiation across mainland and island sites

Can Gio–Dong Nai had the lowest pairwise *F*_*ST*_ under both earlier and primary calls (Table 2, Figure 2). Con Dao–Phu Quoc had the highest point estimate on the full panel (0.457 and 0.405, respectively); on the nine-locus panel its estimates fell to 0.237 and 0.226, and Can Gio–Phu Quoc ranked first. Several bootstrap intervals overlapped. Overall 11-locus *F*_*ST*_ was 0.294 with earlier calls, 0.246 with primary calls, and 0.245 under the 30% rule. After the four loci with high Brookfield estimates were excluded, overall *F*_*ST*_ was 0.287, and Can Gio–Dong Nai and Con Dao–Phu Quoc remained the least and most differentiated pairs on the remaining seven loci (Supplementary Dataset S6).

**Table 2.** Pairwise and overall *F*_*ST*_ estimates for the full and reduced panels.

| **Comparison** | **11 loci, earlier** | **11 loci, primary** | **9 loci, earlier** | **9 loci, primary** |
|---|---|---|---|---|
| Can Gio–Dong Nai | -0.031 | -0.011 | -0.029 | -0.013 |
| Can Gio–Con Dao | 0.343 | 0.314 | 0.268 | 0.248 |
| Can Gio–Phu Quoc | 0.313 | 0.279 | 0.303 | 0.285 |
| Dong Nai–Con Dao | 0.331 | 0.288 | 0.254 | 0.233 |
| Dong Nai–Phu Quoc | 0.291 | 0.261 | 0.267 | 0.255 |
| Con Dao–Phu Quoc | 0.457 | 0.405 | 0.237 | 0.226 |
| Overall | 0.294 | 0.246 | 0.220 | 0.204 |

Negative point estimates are shown as calculated and interpreted as negligible differentiation. Figure 2 gives 95% bootstrap intervals for the primary estimates.

**Figure 2.**
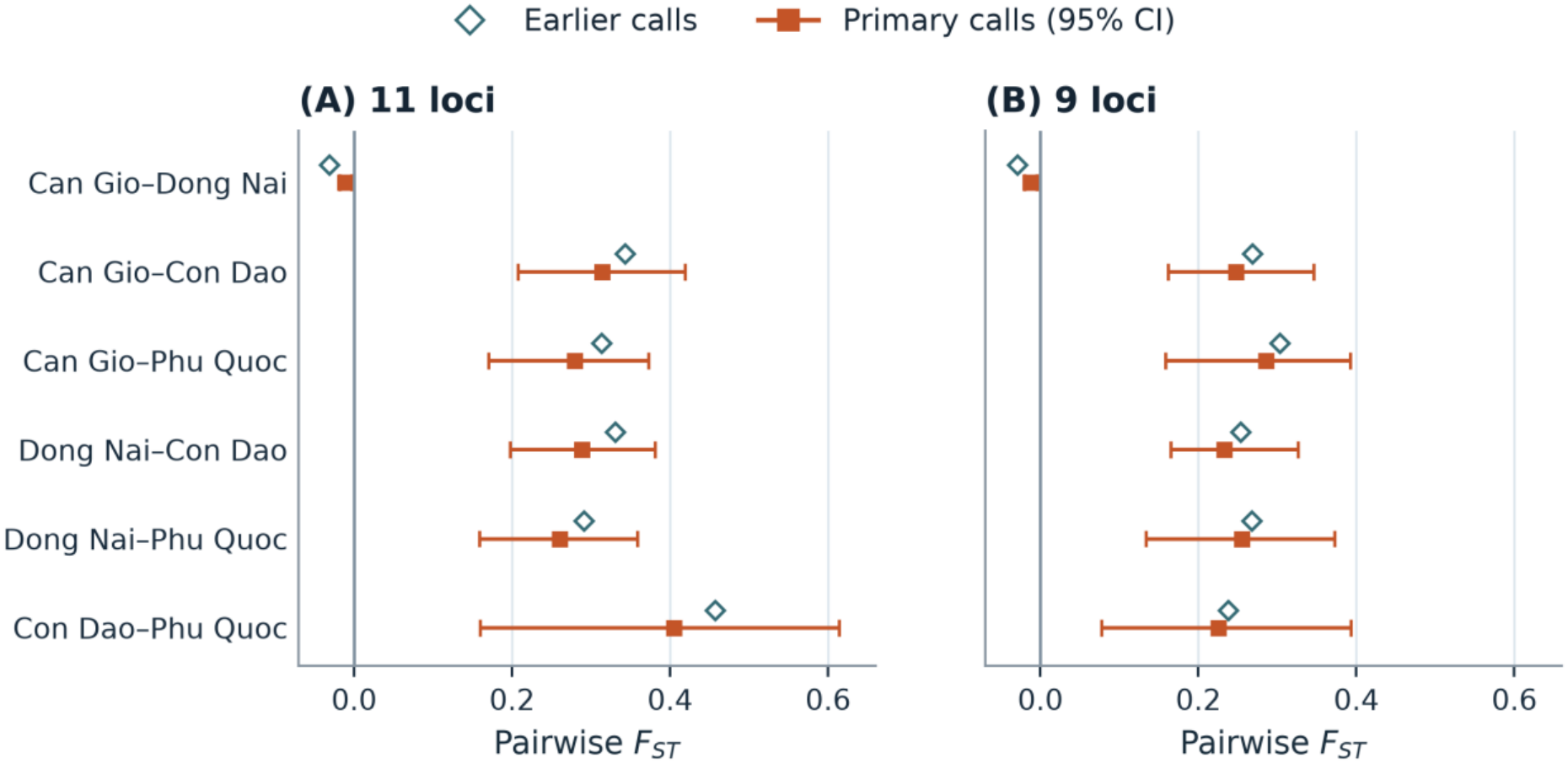
Pairwise *F*_*ST*_ on the 11-locus panel (A) and the nine-locus panel (B). Filled symbols and horizontal bars show primary estimates and 95% bootstrap intervals. Open symbols show estimates from earlier calls.

Restoring the 23 adjacent-peak calls did not change which sites had the highest expected heterozygosity or which pairs were the least and most differentiated (Supplementary Dataset S8). On both panels, the expected pooled capture of either 30-tree allocation changed by less than 0.01 allele, and the joint probability of retaining the two alleles consistently observed only at Con Dao was unchanged.

Most of the molecular variance lay within sites (Table 3): 71.72% on the full panel and 74.06% on the nine-locus panel under primary scoring. The mainland–island grouping explained 12.51% and 18.71%, respectively; its permutation probability of 0.333 is the smallest value possible with only three pairings. The first two axes of the five-locus ordination captured 32.3% of positive eigenvalue variation in *L. littorea* and 29.8% with the congener samples added (Figure 3).

**Table 3.** Hierarchical analysis of molecular variance.

| **Component** | **11 loci, earlier** | **11 loci, primary** | **9 loci, earlier** | **9 loci, primary** |
|---|---|---|---|---|
| Among mainland and island groups | 12.20% | 12.51% | 20.55% | 18.71% |
| Among sites within groups | 21.08% | 15.77% | 7.83% | 7.24% |
| Within sites | 66.72% | 71.72% | 71.62% | 74.06% |
| Overall molecular differentiation | 0.333 | 0.283 | 0.284 | 0.259 |
| Group-level permutation *p* | 0.333 | 0.333 | 0.333 | 0.333 |

The four sites permit three distinct partitions into two groups of two sites.

**Figure 3.**
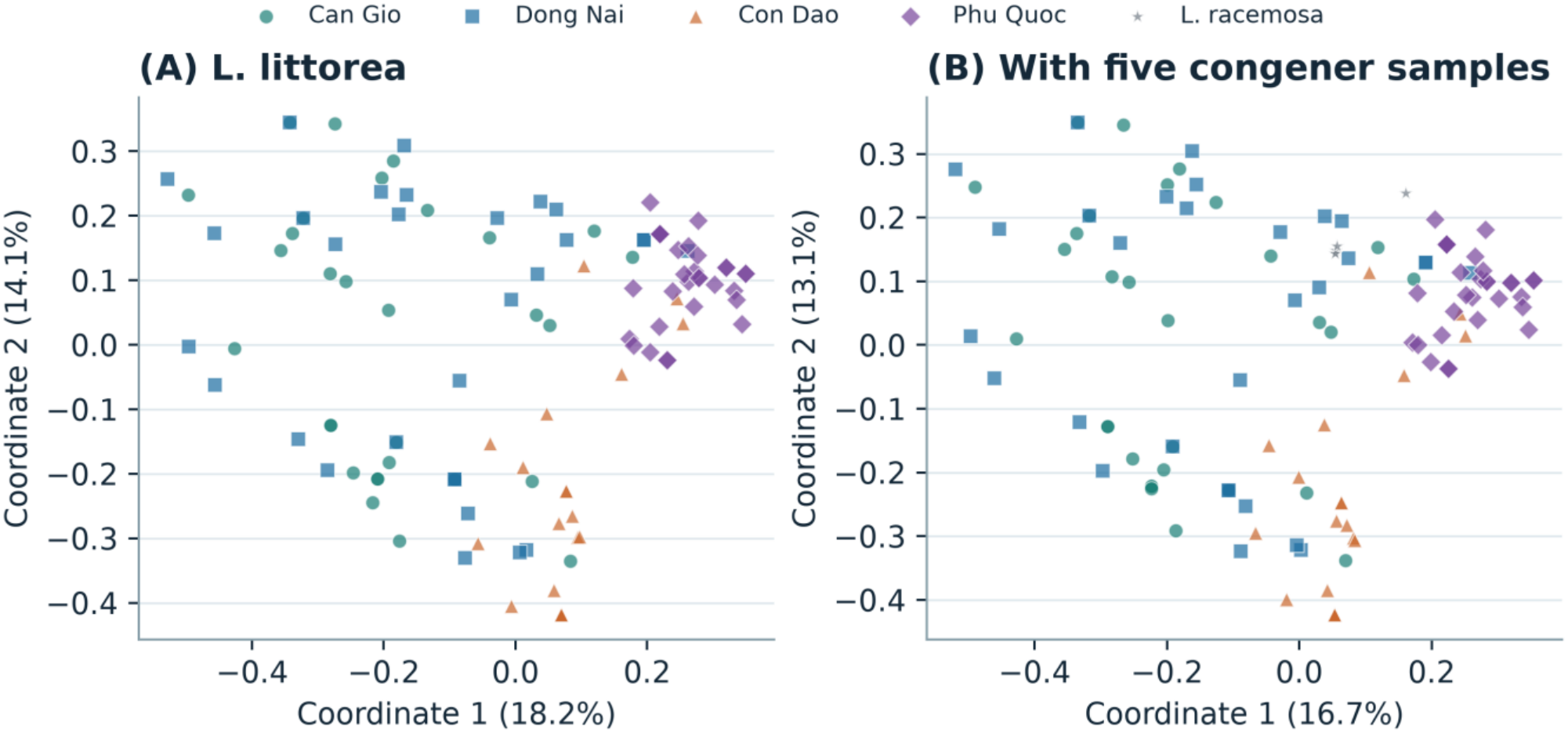
Principal coordinate analysis on the five loci without unresolved calls for 110 *L. littorea* trees (A) and with five *L. racemosa* samples included as interspecific references (B). Overlapping symbols may represent multiple trees.

### 4.3 Allele capture under collection constraints

Of the 5,236 feasible allocations of 30 trees, 286 had at least five trees per source. The observed pool held 37 alleles across 11 loci and 30 across nine loci. Within the five-per-source set, 6–8–11–5 (Can Gio–Dong Nai–Con Dao–Phu Quoc) had the highest minimum relative capture across the compared scenarios, although neighbors such as 5–9–11–5 were practically tied (0.9993 and 0.9992, respectively). With earlier calls, 6–8–11–5 retained an expected 36.73 alleles on the full panel and 29.90 on the reduced panel; with primary calls, it retained 36.89 and 29.94. The nearly even 8–7–8–7 allocation retained 36.61 and 29.76 alleles with earlier calls and 36.81 and 29.86 with primary calls (Table 4). Its two extra Phu Quoc trees raised within-source representation for that site on both panels (Figure 4).

**Table 4.** Allele capture under selected allocations of 30 maternal trees.

| **Allocation** | **Trees** | **11 loci, earlier** | **11 loci, primary** | **9 loci, earlier** | **9 loci, primary** | **All private, earlier** | **All private, primary** |
|---|---|---|---|---|---|---|---|
| At least five per source, highest minimum capture | 6–8–11–5 | 36.73 | 36.89 | 29.90 | 29.94 | 94.8% | 97.2% |
| Nearly even allocation | 8–7–8–7 | 36.61 | 36.81 | 29.76 | 29.86 | 80.0% | 87.5% |
| Unconstrained optimum, 11 loci, earlier calls | 0–17–13–0 | 36.94 | — | — | — | — | — |
| Unconstrained optimum, 11 loci, primary calls (40%) | 0–16–14–0 | — | 36.97 | — | — | — | — |
| Unconstrained optimum, 9 loci, earlier calls | 15–0–15–0 | — | — | 29.99 | — | — | — |
| Unconstrained optimum, 9 loci, primary calls (40%) | 14–0–16–0 | — | — | — | 30.00 | — | — |

Tree counts follow Can Gio–Dong Nai–Con Dao–Phu Quoc. The last two columns show the joint capture of all alleles observed only in Con Dao for the corresponding matrix. In the primary matrix, the apparent restriction of LL-379 allele 264 depends on the unresolved call at C18.

**Figure 4.**
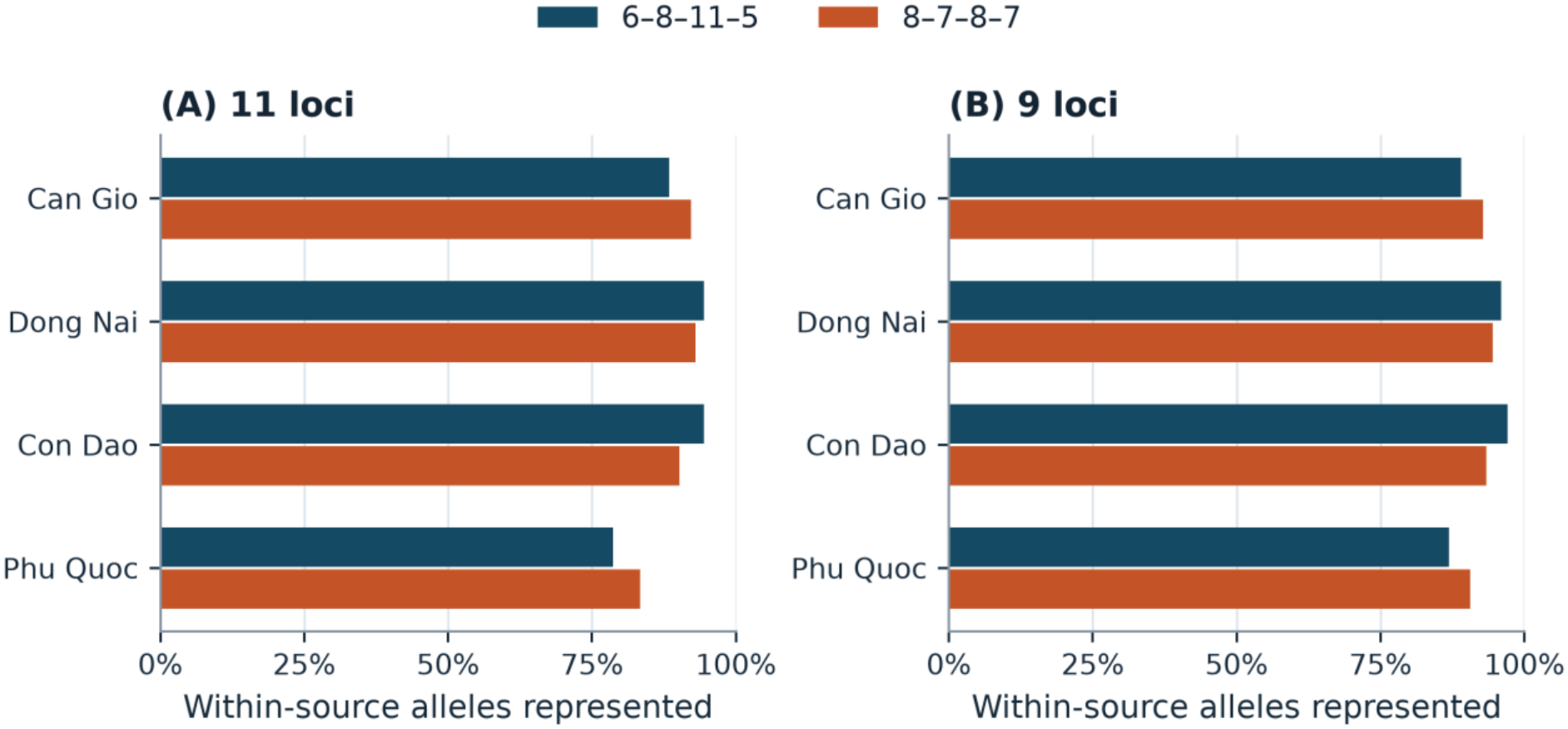
Expected representation of within-source alleles under the two 30-tree allocations with primary scoring, on the 11-locus panel (A) and the nine-locus panel (B). Allocation labels follow Can Gio–Dong Nai–Con Dao–Phu Quoc.

Under primary scoring, three alleles were observed only in the sampled Con Dao trees: 156 at LL-2749 in 11 of 20 trees, 256 at LL-2952 in four, and 264 at LL-379 in six. The first two were confined to Con Dao with or without the 1 bp filter, whereas the apparent restriction of LL-379 allele 264 depended on the unresolved call for Can Gio sample C18 (Supplementary Datasets S2 and S8). The estimated joint probability of keeping all the Con Dao-only alleles was 97.2% for 6–8–11–5 and 87.5% for 8–7–8–7 in the primary matrix, compared with 94.8% and 80.0% with earlier calls (Table 4). For the two consistently Con Dao-only alleles, the primary probabilities were 97.4% and 89.8%. The unconstrained optima left out Phu Quoc and one mainland site. With at least five trees per source, the gain from 30 to 40 trees was smaller than the gain from 20 to 30 (Figure 5).

**Figure 5.**
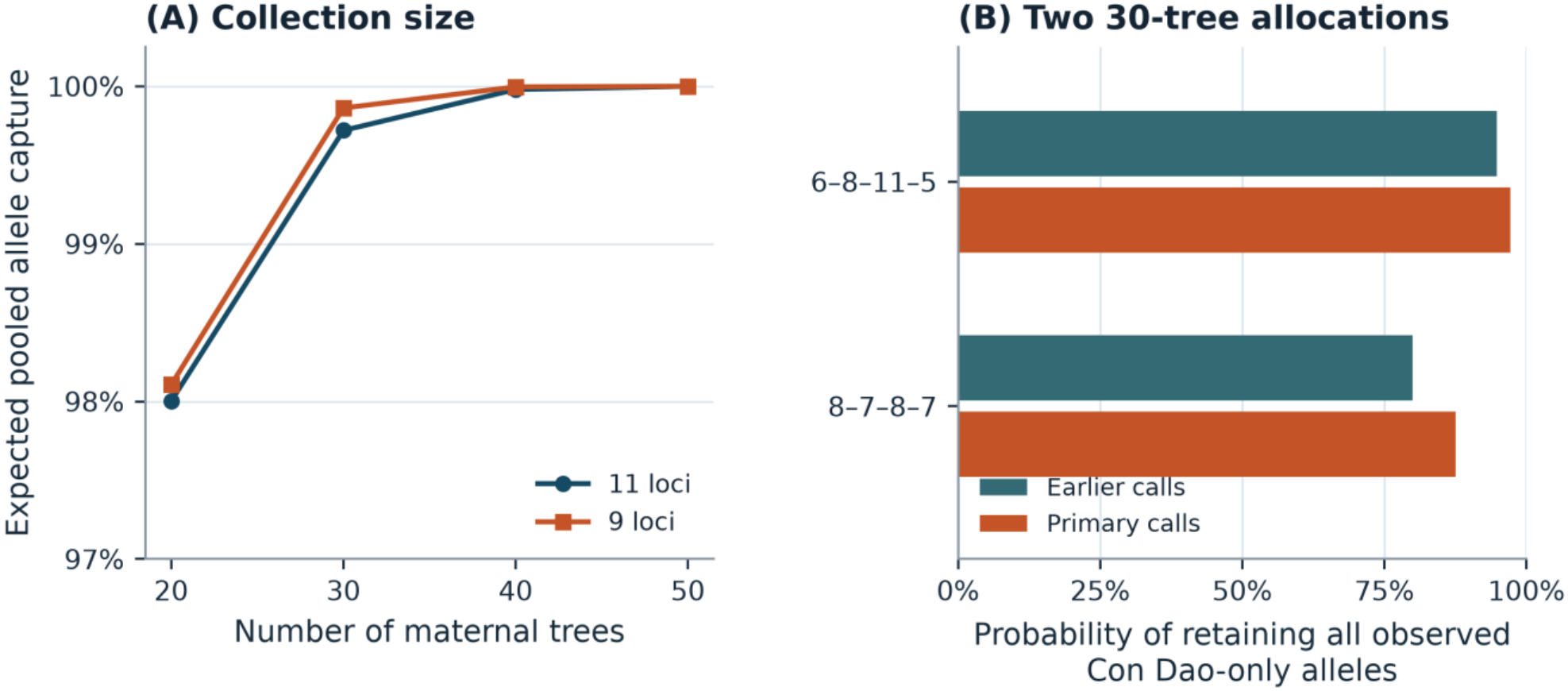
Expected pooled allele capture as the number of maternal trees increases with a minimum of five per source under primary scoring (A), and the joint capture of Con Dao-only alleles under two 30-tree allocations in each scoring matrix (B).

## 5 Discussion

### 5.1 Collection decisions under a fixed budget

Both 30-tree allocations were expected to retain nearly all SSR alleles observed across the four sampled sites. The 6–8–11–5 allocation puts the most trees at Con Dao, the only site with observed private alleles under both earlier and primary calls, and keeps those alleles with a higher joint probability than the even split. The 8–7–8–7 allocation moves two trees to Phu Quoc. That raises the expected representation of Phu Quoc alleles and costs only about 0.12 alleles in pooled capture on the full panel under earlier calls. Neighboring allocations scored almost the same, so we would not read much into the exact counts; what matters is that every source is sampled and Con Dao gets enough trees to hold on to its private alleles. Which allocation to prefer depends on the restoration goal, because keeping an allele somewhere in the collection is not the same as representing the variation within each provenance.

If pooled capture were the only criterion, dropping a source would look attractive, and the unconstrained optima did exactly that, leaving out Phu Quoc and one mainland site. A collection without a site, however, cannot supply seed from that provenance, including any phenotypic or adaptive variation that this marker panel does not detect. Recording the provenance and maternal tree of every seed lot makes it possible to compare survival and growth in the nursery and field, and keeps options open when material is chosen for a particular restoration site. The sampled trees give that decision a quantitative starting point. Work on restored *Rhizophora mucronata* populations elsewhere in Southeast Asia makes the same case for representing existing source variation (Mugundha-Kumar et al., 2026). The small gain in pooled capture beyond 30 trees applies only to the alleles and collection constraints examined here.

### 5.2 Diversity and regional context

After sample sizes were matched, the mainland sites still had the highest expected heterozygosity, as the ranking of ISSR band diversity for these localities had suggested (Huynh et al., 2026). SSR and ISSR estimates measure different features of the marker data, so the agreement is in the ranking, not in the values. Both marker systems also found Can Gio–Dong Nai to be the least differentiated pair. They part ways on the most differentiated pair: Dong Nai–Phu Quoc in the ISSR data, whereas in the SSR data the answer depended on the marker panel. Removing LL-4058 and LL-4164 lowered the Con Dao–Phu Quoc estimate under both scoring matrices, so the size of the contrast between the two islands depends partly on which loci are included.

Guo et al. (2021) reported expected heterozygosities of 0.281–0.444 for five populations of 21–24 trees each. Can Gio and Dong Nai were above that range under both scoring matrices, and Phu Quoc moved from just below it to inside it under primary scoring. Rarefied to 14 allele copies, richness in the primary matrix was 2.617 at Can Gio, 2.749 at Dong Nai, 2.407 at Con Dao, and 2.062 at Phu Quoc, against a median of 2.484 for the 17 richness estimates in Table 2 of Guo et al. (2021). Their rarefaction denominator was not reported, so the median gives only a sense of scale; a joint analysis would need individual multilocus data and allele sizes calibrated across platforms (Morin et al., 2009). Landscape-scale patterns across the Indonesian Archipelago (Manurung et al., 2023) offer a wider biogeographic comparison, but with four sites we have little leverage to separate site history, isolation, and contemporary dispersal.

### 5.3 Limits of interpretation

Heterozygote deficiency was strongest at the mainland sites. It became smaller when the calls were reviewed but persisted at several loci. Subdivision within sites, the mating system, null alleles, and allele-calling uncertainty can all contribute to this pattern. Part of it is locus-specific: at LL-2952 not a single mainland tree was scored as heterozygous, which suggests null alleles or allelic dropout. Heterozygote deficiency persisted after the four loci with high Brookfield estimates were excluded. Can Gio includes several sampling areas (Huynh et al., 2026), and partial self-pollination has been documented in *L. littorea* in China (Zhang et al., 2017). Telling these explanations apart in southern Vietnam would take a hierarchical sampling design and independent evidence on mating or allele transmission. The LL-144 signal outside the scoring window needs product-specific validation before it can enter the genotype matrix. Excluding overlapping adjacent peaks can reduce observed heterozygosity even when missing calls show no detectable concentration by site.

The variance components describe only the four sites we sampled, and with two sites per group there are just three ways to pair them, so group-level inference is coarse. Likewise, the capture probabilities refer to alleles in the sampled maternal trees. What ends up in seeds, seedlings, and established stands can shift through mating, differential fecundity, germination, and survival. The restriction of two observed alleles to Con Dao was robust to the peak-scoring scenarios examined here. Their functional significance is unknown.

### 5.4 Conclusions

Spread across all four sources, 30 maternal trees can retain nearly all of the observed pooled alleles without dropping any provenance. Within that design, a 6–8–11–5 allocation favors the private alleles observed at Con Dao and an 8–7–8–7 allocation gives Phu Quoc fuller representation; this trade-off held across scoring matrices and marker panels. If restoration programs record maternal identity and keep provenances available for follow-up, they can test how these collection choices carry through to seedling representation and establishment.

## Supporting information

Supplementary Datasets S1-S8 and analysis code

## 6 Author contributions

Conceptualization and study design, D.-H.H. and H.-D.T.; investigation, field work, laboratory work, and data curation, T.-K.B.-N. and T.-C.N.; funding acquisition and project administration, D.-H.H.; writing—original draft preparation, D.-H.H.; writing—review and editing, D.-H.H. and H.-D.T.

## 8 Acknowledgments

The authors thank the Department of Science and Technology of Ho Chi Minh City for financial support and the Management Board of Protection and Special-use Forests of Ho Chi Minh City for field coordination and logistical support during sample collection and site verification.

## 10 Data availability statement

The genotype matrices, repeat-peak review, source-file manifest, primer sequences, and the complete 30-tree allocation results are included as Supplementary Datasets S1–S5, and the null-allele sensitivity analysis and site–locus null-allele estimates as Supplementary Datasets S6 and S7. Supplementary Dataset S8 compares the primary results with the matrix that retained the 23 adjacent-peak calls. The accompanying Python scripts reproduce the analyses. The published ISSR comparison is available in Huynh et al. (2026).

## 11 Generative AI statement

OpenAI ChatGPT (Codex) assisted with manuscript language editing, formatting, and writing analysis scripts. The authors reviewed the manuscript and take responsibility for its content.

