## Supplementary Tables S1-S4 for "Allocating maternal-tree collections to conserve genetic diversity of *Lumnitzera littorea* in southern Vietnam"

Supplementary Material

### 1 Supplementary datasets

S1. Genotype matrix for 110 *Lumnitzera littorea* trees and five *L. racemosa* reference samples at 11 loci. The final_, earlier_, and threshold30_ columns give the primary, historical, and 30% calls. The before_1bp_filter_ columns preserve the pre-filter 40% calls. The unresolved_adjacent_peaks flag identifies the 23 excluded combinations.

S2. Peak review for 163 tree–locus combinations, including source pages, peak sizes, intensities, and calls before and after screening for adjacent peaks.

S3. Source-file names, file sizes, and SHA-256 digests.

S4. The 22 primer sequences as supplied for the 11 SSR loci.

S5. Expected allele capture for all 5,236 allocations of 30 maternal trees. The columns specify both marker panels under each of the three scoring matrices.

S6. Pairwise and overall differentiation, sample-size-matched expected heterozygosity, and within-site fixation indices for the full marker panel and the sensitivity analyses that excluded selected loci.

S7. Brookfield estimates of null allele frequency and adjusted Hardy–Weinberg test results for each polymorphic site–locus combination in the primary genotype matrix.

S8. Sensitivity of diversity, differentiation, and allele capture to retaining the 23 calls from overlapping peaks 1 bp apart.

The datasets and analysis code are included in Supplementary_Data.zip. The script recompute_ssr.py reproduces the main analyses, with results provided in SSR_results.json. The scripts null_allele_sensitivity.py and adjacent_peak_sensitivity.py regenerate S6–S8. README_EN.txt gives instructions for all three scripts.

### 2 Supplementary tables

**Supplementary Table S1. Unresolved genotypes by locus in the primary matrix.**

| **Locus** | **Can Gio** | **Dong Nai** | **Con Dao** | **Phu Quoc** | **Total** | **Holm *p*** |
| --- | --- | --- | --- | --- | --- | --- |
| LL-144 | 0 | 0 | 0 | 0 | 0 | — |
| LL-379 | 1 | 0 | 1 | 0 | 2 | 1.000 |
| LL-1086 | 1 | 0 | 4 | 5 | 10 | 0.096 |
| LL-1150 | 4 | 4 | 1 | 1 | 10 | 1.000 |
| LL-4058 | 0 | 1 | 0 | 0 | 1 | 1.000 |
| LL-4164 | 3 | 4 | 1 | 1 | 9 | 1.000 |

**Supplementary Table S2. Sample-size-matched *H_E_* and overall differentiation.**

| **Matrix** | **Loci** | **Can Gio** | **Dong Nai** | **Con Dao** | **Phu Quoc** | **Overall *F_ST_*** |
| --- | --- | --- | --- | --- | --- | --- |
| Earlier scoring | 11 | 0.519 | 0.549 | 0.316 | 0.265 | 0.294 |
| Earlier scoring | 9 | 0.513 | 0.535 | 0.358 | 0.307 | 0.220 |
| Primary scoring (40%) | 11 | 0.524 | 0.554 | 0.340 | 0.296 | 0.246 |
| Primary scoring (40%) | 9 | 0.515 | 0.537 | 0.379 | 0.316 | 0.204 |
| Threshold sensitivity (30%) | 11 | 0.524 | 0.554 | 0.342 | 0.296 | 0.245 |
| Threshold sensitivity (30%) | 9 | 0.516 | 0.537 | 0.382 | 0.316 | 0.203 |

**Supplementary Table S3. Summary of Hardy–Weinberg tests.**

| **Matrix** | **Site** | **Polymorphic combinations** | **Two-sided *q* < 0.05** | **Heterozygote deficit *q* < 0.05** |
| --- | --- | --- | --- | --- |
| 40% primary calls | Can Gio | 10 | 9 | 9 |
| 40% primary calls | Dong Nai | 10 | 6 | 5 |
| 40% primary calls | Con Dao | 9 | 5 | 0 |
| 40% primary calls | Phu Quoc | 8 | 4 | 3 |
| 30% sensitivity | Can Gio | 10 | 9 | 9 |
| 30% sensitivity | Dong Nai | 10 | 5 | 5 |
| 30% sensitivity | Con Dao | 9 | 5 | 0 |
| 30% sensitivity | Phu Quoc | 8 | 4 | 3 |

**Supplementary Table S4. Sampling budgets with primary scoring.**

| **Loci** | **Trees** | **Feasible designs with minimum five per site** | **Best allocation** | **Expected pooled alleles** | **Observed pooled alleles** |
| --- | --- | --- | --- | --- | --- |
| 11 | 20 | 1 | 5–5–5–5 | 36.26 | 37 |
| 11 | 30 | 286 | 5–10–10–5 | 36.90 | 37 |
| 11 | 40 | 1736 | 5–16–14–5 | 36.99 | 37 |
| 11 | 50 | 4671 | 5–23–17–5 | 37.00 | 37 |
| 9 | 20 | 1 | 5–5–5–5 | 29.43 | 30 |
| 9 | 30 | 286 | 8–5–12–5 | 29.96 | 30 |
| 9 | 40 | 1736 | 14–5–16–5 | 30.00 | 30 |
| 9 | 50 | 4671 | 23–5–17–5 | 30.00 | 30 |
